# Multimodal Optical Imaging and Modulation through Smart Dura in Non-Human Primates

**DOI:** 10.1101/2025.02.27.640384

**Authors:** Nari Hong, Sergio Montalvo Vargo, Gaku Hatanaka, Zhaoyu Gong, Noah Stanis, Jasmine Zhou, Tiphaine Belloir, Ruikang K. Wang, Wyeth Bair, Maysamreza Chamanzar, Azadeh Yazdan-Shahmorad

**Affiliations:** Department of Bioengineering, University of Washington, Seattle, WA, 98195, USA; Washington National Primate Research Center, Seattle, WA, 98195, USA; Department of Electrical and Computer Engineering, Carnegie Mellon University, Pittsburgh, PA, 15213, USA; Department of Neurobiology and Biophysics, University of Washington, Seattle, WA, 98195, USA; Department of Biomedical Engineering, Carnegie Mellon University, Pittsburgh, PA, 15213, USA; Carnegie Mellon Neuroscience Institute, Pittsburgh, PA, 15213, USA; Department of Electrical and Computer Engineering, University of Washington, Seattle, WA, 98195, USA; Weill Neurohub

## Abstract

Multimodal neural interfaces that integrate electrical and optical functionalities are promising tools for neuroscientific and clinical applications that involve recording and manipulating neuronal activity. However, most technologies for multimodal implementation are largely restricted to small animal models and lack the ability to translate to the larger brains of non-human primates (NHPs). Smart Dura, a recently developed large-scale neural interface for NHPs, enables high-density electrophysiological recordings and broad optical accessibility, providing multiscale information with enhanced spatiotemporal resolution. In this paper, the multimodal capabilities of Smart Dura are demonstrated through integration with multiphoton imaging, optical coherence tomography angiography (OCTA), and intrinsic signal optical imaging (ISOI), as well as optical manipulations such as photothrombotic lesioning and optogenetics. Through the transparent Smart Dura, in vivo fluorescence vascular imaging is achieved down to depths of 200 and 550 μm using two-photon and three-photon microscopy, respectively. When combined with simultaneous electrophysiology, Smart Dura also enables assessment of vascular and neural dynamics via OCTA and ISOI, the induction of ischemic stroke, and the application of optogenetic neuromodulation across a wide cortical area of 20 mm in diameter. These capabilities support comprehensive investigations of brain dynamics in NHPs, advancing translational neurotechnology for human applications.

## 1. Introduction

Mapping the functional structure of the brain is a long-standing challenge in neuroscience research. Various neurotechnologies to study neural circuits at multiple spatial and temporal scales have been developed to record and manipulate neuronal activity, based on two main modalities: electrical and optical.^[1]^ Electrical interfaces, such as penetrating probes and electrocorticography (ECoG), are capable of electrophysiological recording and stimulation with high temporal resolution but lack spatial precision or completeness of coverage. On the other hand, optical imaging techniques, such as fluorescence calcium imaging and intrinsic signal imaging, provide high spatial resolution for structural and functional imaging across continuous regions but have drawbacks, including low temporal resolution or insufficient signal-to-noise ratio. In addition, optical manipulation techniques offer several advantages over electrical stimulation. For instance, optogenetics facilitates cell-type-specific neuromodulation with high spatial and temporal resolution and allows for artifact-free recording.^[2]^ Combining electrical and optical approaches present unique opportunities to investigate both brain function and dysfunction. A multimodal interface that enables simultaneous electrical and optical interfacing with the brain and leverages the strengths of each modality can serve as a powerful tool for integrating complementary measurements and manipulations with unprecedented spatiotemporal resolution.

Recent advances in implementing multimodal integration have focused on developing flexible and transparent electrophysiological neural interfaces and combining these with various optical imaging and manipulation paradigms.^[3–5]^ To fabricate multimodal interfaces, flexible polymers such as polyimide, polyethylene terephthalate (PET), Polyethylene naphthalate (PEN), Parylene C, and polydimethylsiloxane (PDMS) have been used as substrates. Moreover, transparent electrodes have been realized for further enhanced optical transparency by employing materials such as ultrathin gold, graphene, indium tin oxide (ITO), carbon nanotubes, and PEDOT:PSS. Using the multimodal interfaces, previous studies have demonstrated intracortical^[6,7]^ or surface^[8–11]^ recordings combined with wide-field or two-photon calcium imaging. In addition, simultaneous electrophysiological recording and optical imaging have been attempted in conjunction with voltage-sensitive dye imaging^[12]^ and intrinsic signal imaging.^[13]^ Furthermore, the application of optogenetic modulation and the recording of the corresponding electrophysiological responses have also been demonstrated.^[10,14–17]^ Despite these significant recent efforts, most techniques are still limited in their applications to small animal models, such as rodents. To ultimately be applicable and translatable to humans, the interface needs to be scaled up to allow recording and imaging over wider coverage areas as well as neuromodulation across larger volumes. Moreover, this needs to be validated in larger brains, such as non-human primates (NHPs), as they can bring relevant translational insights given their high cognitive and behavioral complexity.^[18,19]^

We recently developed a large-scale multimodal neural interface for NHPs, called Smart Dura,^[20,21]^ to address this outstanding need. Smart Dura is a functional version of an artificial dura^[22–25]^ that can replace the native dura, providing electrophysiological recordings and high optical accessibility to large cortical areas. Following our previous work focusing on the design and microfabrication of the device as well as benchtop and in vivo validation, in this study, we present the multimodal functionality of Smart Dura for optical imaging, perturbation, and manipulation applications with simultaneous electrophysiology (**Figure 1a**). We demonstrate the feasibility of combining Smart Dura with multiphoton imaging, optical coherence tomography angiography (OCTA), and wide-field intrinsic signal optical imaging (ISOI), which are utilized to assess neural and vascular functions. We also show that two optical manipulations for modulating vascular and neural dynamics, photothrombotic lesioning and optogenetic stimulation, can be applied to the brain through Smart Dura during neural recording.

**Figure 1.**
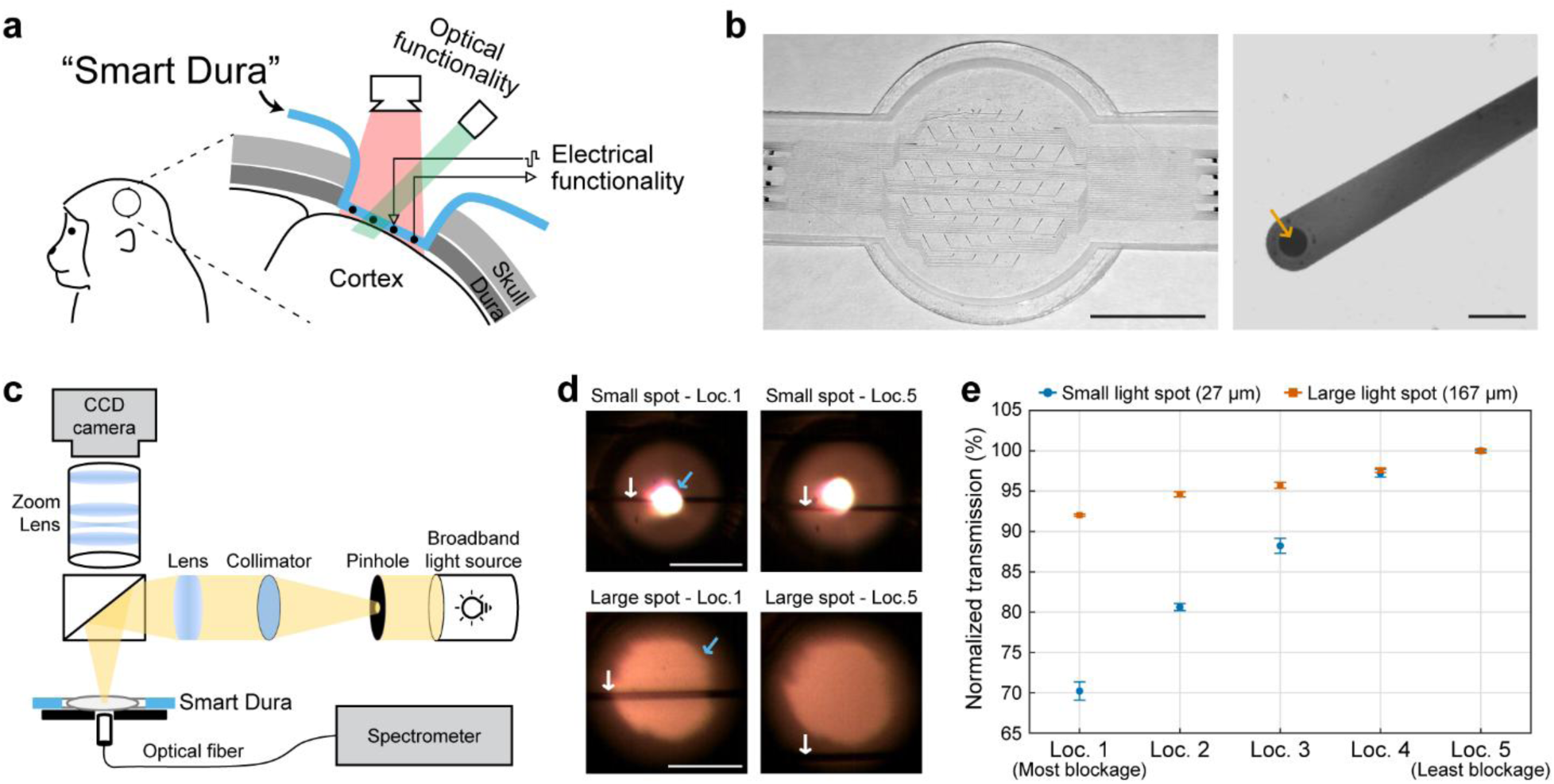
Smart Dura as a large-scale and transparent multimodal neural interface. (a) Schematic illustration of the electrical and optical functionalities of Smart Dura. (b) Photograph of the 64-channel Smart Dura and micrograph of the metal trace and electrode. The orange arrow indicates the PEDOT:PSS-coated microelectrode. Scale bar: 10 mm and 50 μm. (c) Schematic of the optical setup for high-spatial-resolution transmission measurements. (d) Micrographs of the small and large light spots (27 and 167 μm in diameter, respectively) when they are most (Loc. 1) and least (Loc. 5) blocked by a 10 μm-wide metal trace. The white and blue arrows point out metal traces and light spots, respectively. The exposure time was set long enough to saturate the images to show the metal traces and light spots. Scale bars: 100 μm. (e) Average transmission in the wavelength range of 400-1000 nm for two light spot sizes at five different locations with different light blockage levels, normalized to the value of the least blockage (Loc. 5). Data are presented as the mean ± standard error of the mean. n = 5 measurements.

## 2. Results

### Semi-transparent Smart Dura for Multimodal Applications

Our Smart Dura^[20,21]^ is a large-scale functional artificial dura made of a transparent, flexible, and soft PDMS and Parylene C bilayer substrate that offers clear optical access to the brain with mechanical compliance with brain tissue and minimal invasiveness (Figure 1b). It has 64 microelectrodes fabricated with Pt/Au/Pt stacking and PEDOT:PSS electroplating over a 20 mm diameter (area: 315 mm^2^), enabling electrical recording and stimulation over large cortical regions. Each electrode has a diameter of 20 μm (orange arrow in Figure 1b), facilitating highly localized measurements of high-frequency activity,^[26]^ and is individually connected to a backend connector pad via a 10 μm-wide metal trace for electrical signal transmission. These micron-sized geometries of the electrodes and traces provide enhanced optical transparency (the opaque metal area covers only 2.81% of the total area).

To demonstrate the enhanced optical access of Smart Dura for multimodal applications, we conducted high-spatial-resolution transmission measurements in the wavelength range of 400 nm to 1000 nm, which is the prevalent range used for optical techniques in neuroscience. As shown in the schematic of Figure 1c, we implemented an optical setup to obtain a focused beam of light with a spot size as small as 27 μm in diameter, comparable to the cell body of neurons. We then measured the transmission through the Smart Dura for two light spot sizes of 27 and 167 μm in diameter at five different locations around our 10 μm-wide metal trace. Each location has a different level of light blockage by the metal trace, with Loc. 1 and 5 having the highest and lowest levels of light obstruction, respectively (Figure 1d). Figure 1e shows the average transmission in the wavelength range of 400-1000 nm, normalized to the value of the least blockage (Loc. 5). At the highest blockage level when the trace was aligned to the center of the 27 μm diameter spot (Small spot – Loc. 1 in Figure 1d), the normalized average transmission was 70.22% and it increased as the degree of blocking by the trace reduced (blue circles in Figure 1e and Figure S1a). For the 167 μm diameter spot, the normalized transmission was 92.03% at the most blockage (Large spot – Loc. 1 in Figure 1d), which was less affected by light blockage from the metal trace compared to the smaller spot (orange squares in Figure 1e and Figure S1b). These high levels of transmission, despite being blocked by the opaque metal trace, suggest that a significant amount of light can still pass around the micron-sized traces over the applicable wavelength ranges for multimodal applications.

**Figure S1.**
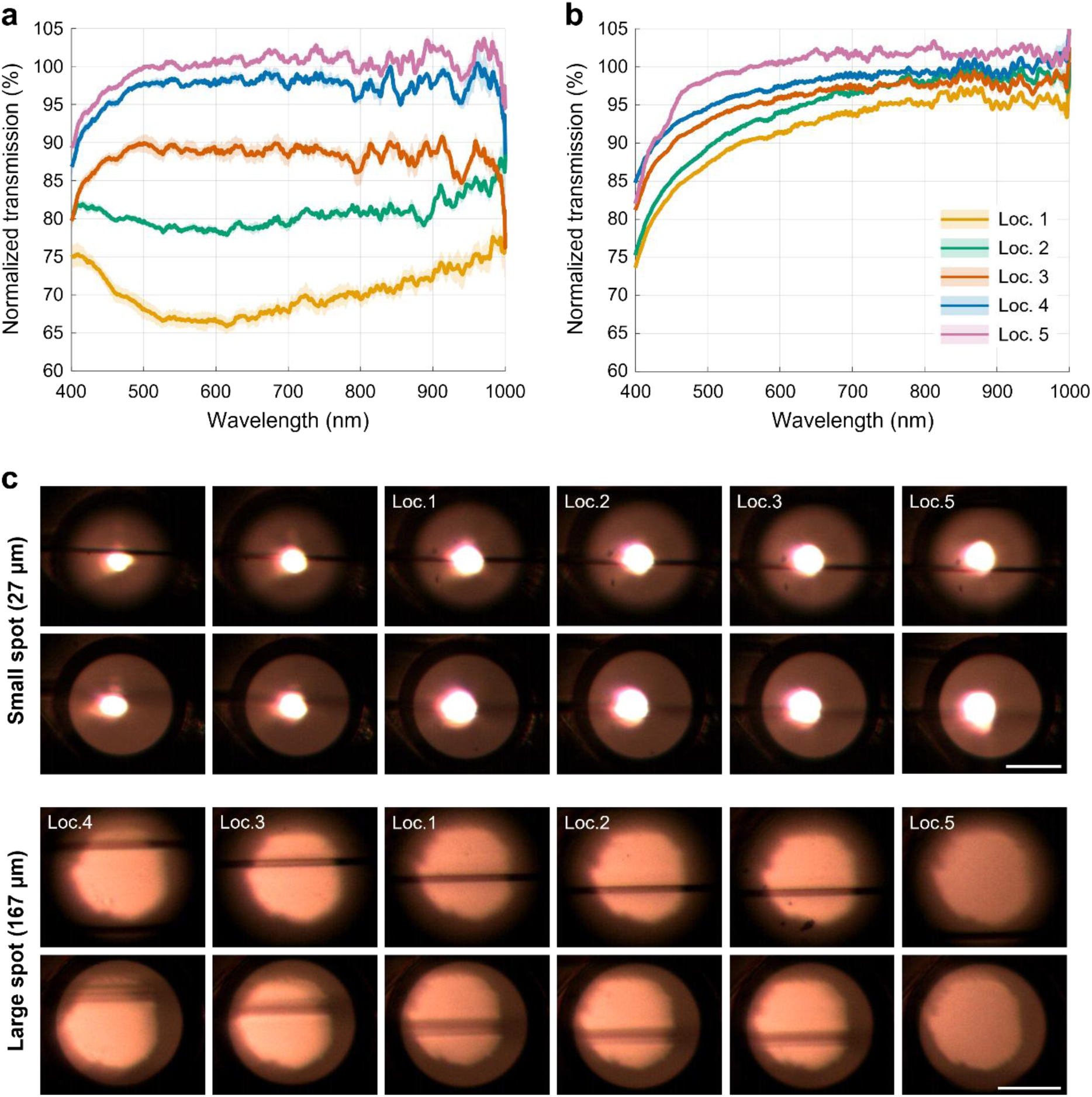
Transmission measurements through Smart Dura with different light blockage levels. (a) Transmission spectra for a 27 μm diameter spot at five locations of a 10 μm-wide metal trace with different light blockage levels. Loc. 1 and Loc. 5 are the locations with the highest and lowest levels of the blockage, respectively. Data are normalized to the average value of the least blockage (Loc. 5) and presented as the mean (solid line) ± standard deviation (shaded area). n = 5 measurements. (b) Transmission spectra for a 167 μm diameter spot at five locations of a 10 μm-wide metal trace with different light blockage levels. (c) 1D-scan transmission measurement for two different spot sizes. For each spot size, the top and bottom rows show images focused on the metal trace and an optical fiber, respectively. The exposure time was set long enough to saturate the images to show the metal traces and light spots. Scale bars: 100 μm.

### 2.2. Multiphoton Imaging through Smart Dura

Multiphoton microscopy has been widely used for structural and functional brain imaging at various depths in many species, including NHPs.^[27,28]^ To validate the optical imaging capability through the transparent Smart Dura and to see how the opaque portion affects imaging at depth, we performed multiphoton imaging both bench-side and in vivo (**Figure 2a**). Using two-photon microscopy with 920 nm excitation light, we first imaged a sample containing fluorescent pollen grains with diameters on the order of ten micrometers, similar to the cell body of neurons, placed under the Smart Dura (Figure 2b). The Smart Dura electrodes were facing downward toward the fluorescent sample, mimicking the geometry in the case of brain imaging with simultaneous electrical recording. As shown in a 3D reconstructed image (left panel in Figure 2b), fluorescence from the pollen grains was successfully detected through the transparent layers of the Smart Dura. Moreover, grains below opaque metal traces could also be detected, supporting the message articulated in the previous section. In the images at the depths of 0 and 219 μm (middle and right panels in Figure 2b), a shadow was observed under the widest part of the metal trace (50 μm across, blue arrow) at the site of the electrode opening, however, this loss of fluorescence intensity was almost negligible below the narrow part of the trace, which is only10 μm wide (orange arrow).

**Figure 2.**
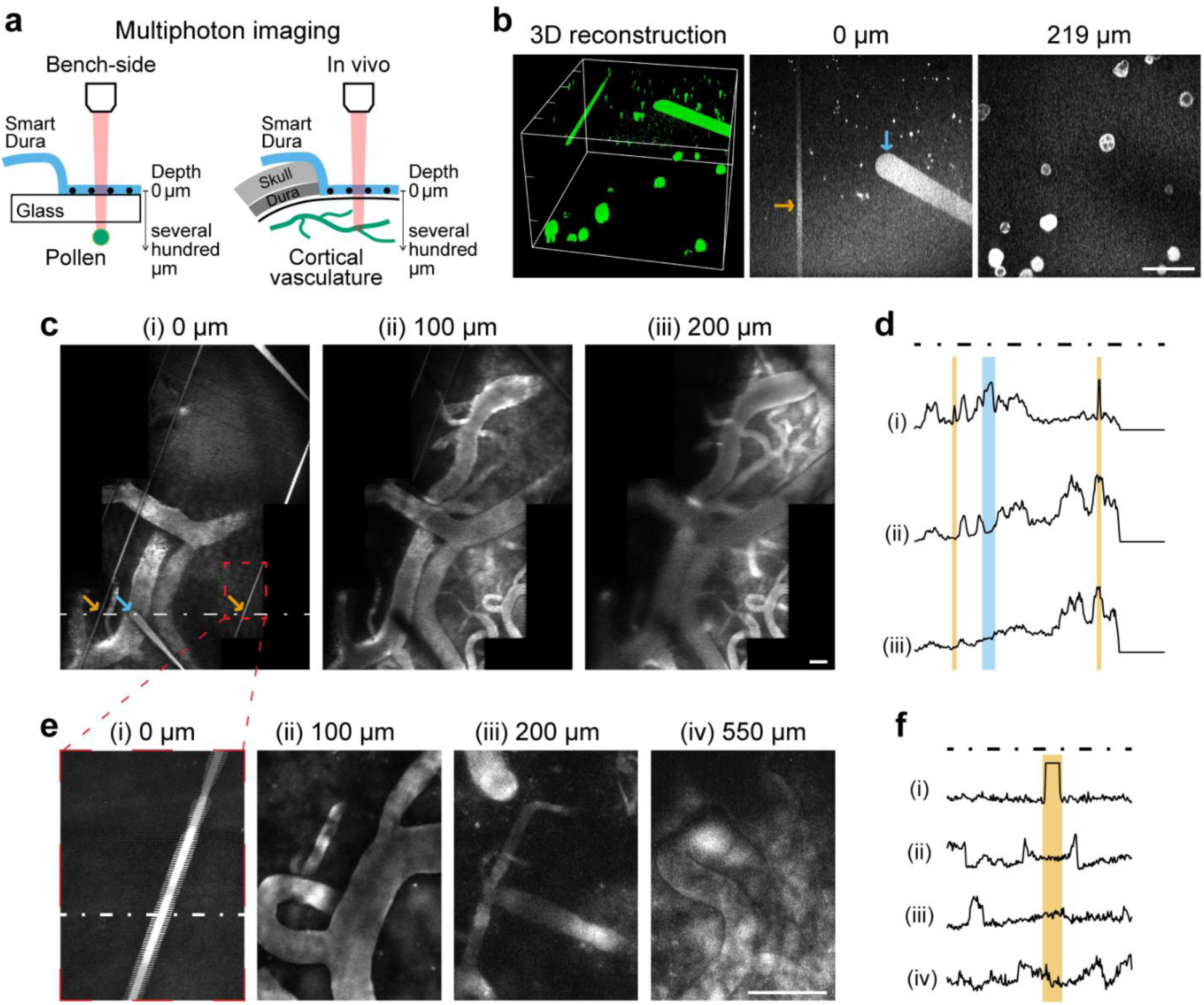
Multiphoton imaging through the Smart Dura. (a) Schematics of bench-side and in vivo multiphoton imaging of pollen and cortical vasculature. The Smart Dura electrodes are facing downward toward the fluorescent sample, which is the same condition as when recording neuronal signals from the brain. The image at a depth of 0 µm is the plane focused on the metal traces. (b) Two-photon imaging of pollen grains placed under the Smart Dura. 3D reconstruction of the two-photon images (left panel) and images at depths of 0 (middle panel) and 219 µm (right panel). The orange and blue arrows in the middle panel indicate metal traces with widths of 10 and 50 μm, respectively. (c) In vivo vascular images at depths of 0, 100, and 200 µm in V1 under the Smart Dura using two-photon microscopy. The orange and blue arrows in the left panel point out metal traces with widths of 10 and 50 μm, respectively. (d) Intensity profiles at depths of 0, 100, and 200 µm (from top to bottom) along the white dashed line shown in the image of 0 µm. The areas highlighted in orange and blue represent the regions where the metal traces of 10 and 50 μm width pass through, respectively. (e) Three-photon images of the vasculature under the Smart Dura. The images were taken from the area highlighted in the red dashed box in Figure 2c. (f) Intensity profiles at different depths (from top to bottom: 0, 100, 200, and 550 µm depth) along the white dashed line shown in the image of 0 µm. The orange shaded area is the region where the 10 μm-wide metal trace passes through. Ticks in the 3D images and scale bars in 2D images: 100 µm.

Next, we conducted in vivo multiphoton imaging of cortical vasculature through the Smart Dura using intravenously administered FITC-dextran in an anesthetized animal. With two-photon microscopy, we could obtain fluorescence images of microvasculature in the primary visual cortex (V1) through the Smart Dura at depths of up to 200 μm (Figure 2c). On the surface, i.e., at a depth of 0 μm (left panel in Figure 2c), the imaging reveals metal traces of the Smart Dura that vary in width from 50 μm taper at the electrode contact (blue arrow) to 10 μm (orange arrows). As shown in the intensity profiles in Figure 2d, the widest trace (50 μm) resulted in a significant intensity reduction at 100 μm depth, but only a modest decrease at 200 μm depth (blue shaded area). On the other hand, the 10 μm-wide trace had little effect on the intensity at both depths of 100 and 200 μm (orange shaded area). We next acquired vascular images using three-photon microscopy with 1300 nm excitation light, which is used to image deeper brain regions,^[29,30]^ and were able to image microvasculature at depths of up to 550 μm (Figure 2e). The metal trace of 10 μm casts negligible shadows on images of the underlying vasculature at depths of 100 μm and deeper. The intensity profiles in Figure 2f also show no noticeable intensity loss caused by the metal trace. These results demonstrate that the Smart Dura is applicable to brain imaging via multiphoton microscopy, as it can image fluorescent targets as small as individual neurons under the Smart Dura, and the narrow traces become effectively transparent for the images beyond a certain depth with insignificant intensity loss. By combining large-scale surface recordings provided by our Smart Dura with multiphoton functional imaging, such as calcium imaging, which captures the activity of single neurons at depths of hundreds of micrometers, it becomes possible to elucidate the relationship between local cellular dynamics in deeper brain regions and global brain state.

### 2.3. Photochemical Ischemic Lesioning and Optical Coherence Tomography Angiography combined with electrical recording and stimulation of Smart Dura

Brain perturbation techniques with simultaneous monitoring of neural activity enable the investigation of brain function and dysfunction. Our Smart Dura with high-density microelectrodes and a large surface area provides high-resolution electrophysiology across a broad cortical region as well as spatially localized electrical stimulation, allowing us to examine the functional impact of perturbations. Moreover, because of its high level of transparency, optical perturbation techniques can be readily applied to Smart Dura. To show this capability, we employed photothrombotic lesioning in combination with electrophysiological recording using Smart Dura. The photothrombotic technique, developed in our recent publication,^[31–35]^ can generate focal ischemic stroke in the NHP cortex photochemically using the photosensitive dye Rose Bengal. To image changes in the vasculature, we performed OCTA before and after stroke induction using a custom-built OCT system.^[36,37]^ We first induced an ischemic lesion in V1 by intravenous administration of Rose Bengal and localized light illumination (**Figure 3a**). OCTA images acquired before and after stroke induction (Figure 3b) show a significant reduction in blood flow centered on the illuminated area. The lesion detected on the OCTA image was approximately 6.5 mm in diameter, consistent with our previous study.^[31]^ We then analyzed stroke-induced changes in neural activity using electrophysiological recordings from Smart Dura. Figure 3c presents gamma band (30-59 Hz) filtered traces from an electrode within the lesion, showing a great decline in amplitude after stroke. As shown in the heatmaps of Figure 3d, the gamma power in the stroke-induced region detected in OCTA images (red dashed circle) was significantly reduced to nearly 0 (p < 0.0001; Paired t-test; n = 12 electrodes), along with the power decrease in the surrounding regions (p < 0.0001; Paired t-test; n = 40 electrodes), similar to our previous studies.^[31,33]^

**Figure 3.**
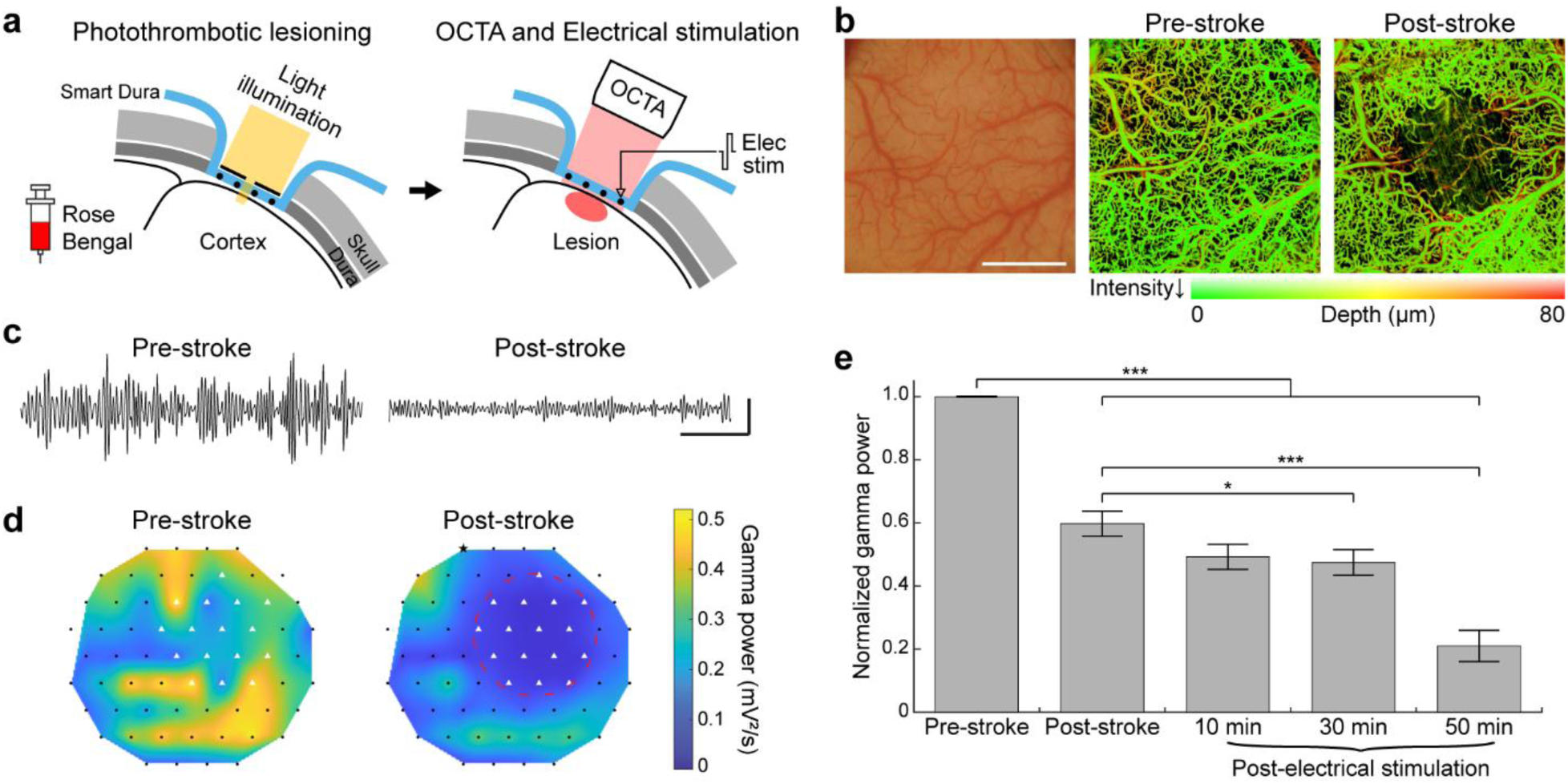
Photochemical ischemic lesioning and optical coherence tomography angiography combined with electrical recording and stimulation of the Smart Dura. (a) Schematics of photothrombotic lesioning through the Smart Dura to induce an ischemic stroke, optical coherence tomography angiography (OCTA) imaging, and neuroprotective electrical stimulation after stroke. (b) The surface of the V1, covered with the Smart Dura (left panel). Ischemic stroke was photochemically induced by illuminating a light in the center of the area. OCTA images before (center panel) and after stroke (right panel), showing the absence of blood flow in the stroke-induced region. Scale bar: 5 mm. (c) Representative gamma band traces of an electrode in the stroke-induced region before and after the stroke. Scale bar: 500 ms and 50 µV. (d) Heatmaps of gamma power before and after the stroke, showing the power decrease in the stroke-induced region detected in OCTA images. A red dashed circle and white triangles denote the stroke-induced region and the electrodes within the area, respectively. (e) Normalized gamma power of electrodes, not in the stroke-induced region, before and after stroke, and after electrical stimulation. Electrical stimuli were delivered to the electrode marked with a star in Figure 3d. Data are presented as the mean ± standard error of the mean. Repeated measures ANOVA with Bonferroni’s post-hoc test. *p < 0.05 and ***p < 0.001. n = 40 electrodes.

Next, to demonstrate that electrical stimulation can be combined with this optical perturbation and electrophysiology, we replicated our recent work on post-stroke acute stimulation for neuroprotection.^[33]^ In this previous study, we have demonstrated that electrical stimulation applied in the early stage of acute ischemic stroke can alleviate excessive depolarization and neuroinflammatory responses surrounding the ischemic core, thereby reducing neuronal death during the ischemic cascade. One of the major physiological findings was that in electrically stimulated animals, the gamma power in the perilesional area gradually decreased, whereas peri-infarct hyperactivation was seen at 1-2 hours after stroke in unstimulated control animals. Here, we delivered theta-burst electrical stimulation 60 minutes after stroke induction. Figure 3e shows the normalized gamma power of the electrodes outside of the lesion in pre-stroke, post-stroke, and post-electrical stimulation periods. Consistent with our previous findings, a progressive gamma power decrease in the peri-infarct region was observed as electrical stimulation continued following the stroke (Figure 3e). Taken together, the results suggest that Smart Dura can be integrated with optical lesioning and imaging to induce acute stroke, monitor neurological and vascular changes, and provide therapeutic electrical stimulation. Notably, the high electrode density of Smart Dura enabled detection of activity changes caused by stroke induction and subsequent electrical stimulation with higher spatial resolution compared to previous studies using 32-channel electrode arrays.^[31,33]^ Such ability holds promise for the development of advanced therapeutic strategies and their successful translation into clinical applications.

### 2.4. Wide-field Intrinsic Signal Optical Imaging with Simultaneous Large-scale Electrophysiology

Integrating electrical recording and optical imaging of neural activity provides a powerful approach to interrogate neural circuits with enhanced spatial and temporal resolution. In particular, large-area electrophysiology combined with wide-field imaging enables the investigation of the functional organization of the large NHP cortices, which is structured at scales spanning from submillimeter to millimeter. Our Smart Dura, which can acquire electrical signals over a large coverage area while at the same time having optical accessibility to a wide cortical area, is well-suited in this regard. To assess the feasibility of simultaneous large-scale electrophysiological recording and optical functional imaging using our Smart Dura, we used wide-field ISOI of hemodynamics, which is an imaging technique to infer neural activity indirectly based on hemodynamic response changes.^[38]^ We imaged evoked responses in V1 to visual stimulation in an anesthetized animal while recording neural signals from the same cortical area using the Smart Dura (**Figure 4a**). Visual stimuli consisting of chromatic and achromatic oriented, drifting square-wave gratings of duration 4 seconds were randomly interleaved and presented separately to the left and right eyes. Figure 4b shows representative spectrograms of neural activity at an electrode during chromatic or achromatic visual stimulation. For both stimuli, signal power increased as the stimuli were applied and such power increases were particularly prominent in the high-gamma (HG) band of 60-150 Hz. We were also able to create two basic functional maps over a 20 mm diameter cortical area with high spatial resolution using intrinsic signals acquired using ISOI: maps for color domains (Figure 4c) and ocular dominance (Figure 4d). Color blobs and ocular dominance columns, which are well-known characteristic features of V1 in macaques,^[39,40]^ were observed in the color map and ocular dominance map, respectively.

**Figure 4.**
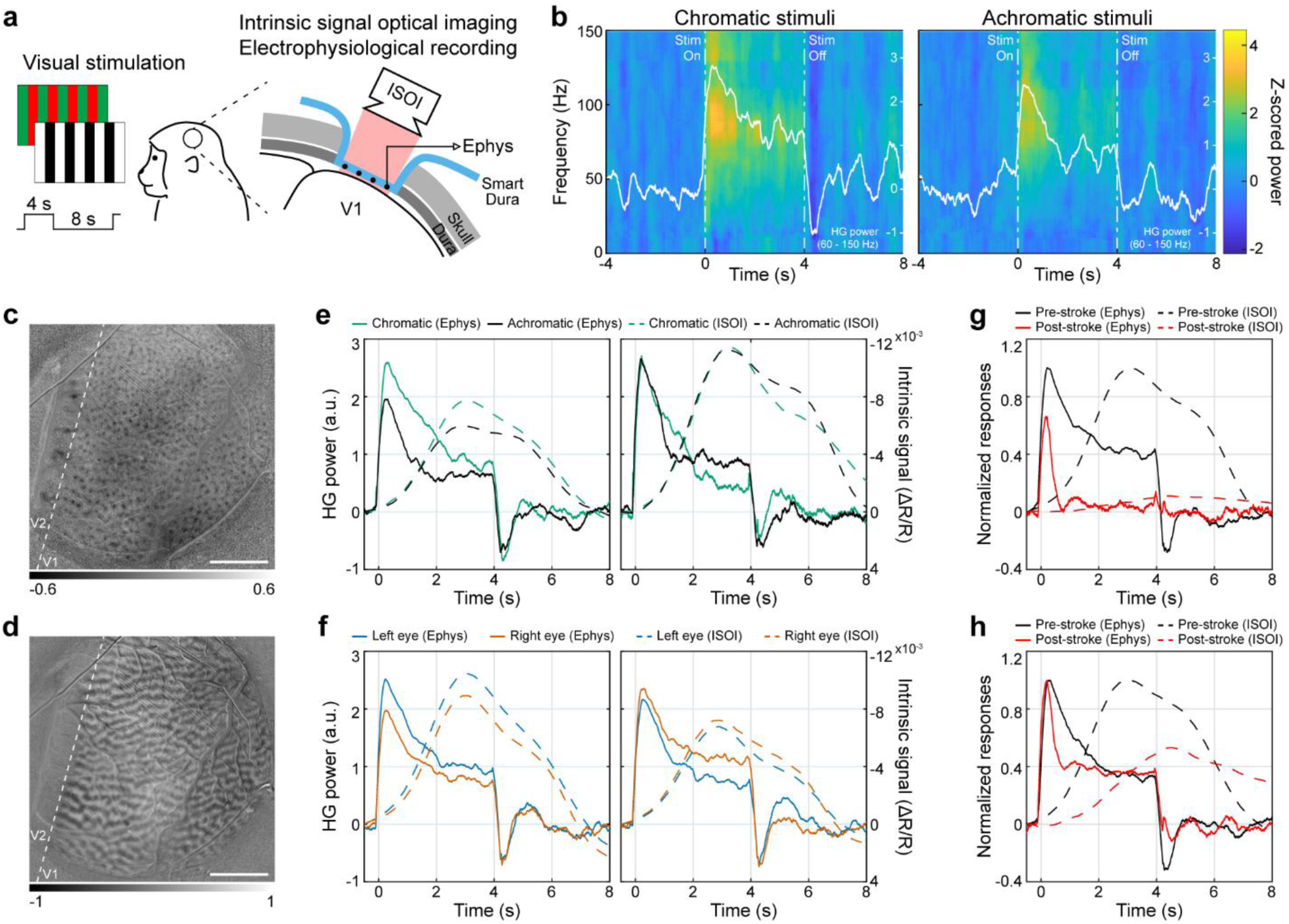
Wide-field intrinsic signal optical imaging through the Smart Dura with simultaneous electrophysiology. (a) Schematics of visual stimulation to the anesthetized animal and simultaneous wide-field intrinsic signal optical imaging (ISOI) and electrophysiological recording (Ephys) from V1. (b) Representative spectrograms for an electrode, showing trial-averaged neural responses during chromatic (left panel) or achromatic (right panel) visual stimulation (10 trials each). The averaged high-gamma (HG) power in the 60-150 Hz frequency band is overlaid in white. The stimulation onset occurred at 0 s and lasted 4 seconds. (c) Color map for left eye stimulation, exhibiting color blobs (dark pixels) that are color-preferring neuronal clusters, and (d) Ocular dominance map, showing ocular dominance columns (dark pixels: right eye; light pixels: left eye). The white dashed line shows the primary and secondary visual cortex border (V1/V2). Scale bar: 5 mm. (e) Trial-averaged HG power via Ephys and intrinsic signal via ISOI in response to chromatic or achromatic stimuli (80 trials each). The left and right panels show the responses of electrodes with the greatest preference for chromatic or achromatic stimuli, respectively. Intrinsic hemodynamic responses were obtained from five pixels near each electrode. The y-axis for intrinsic signals is drawn in reverse. (f) Trial-averaged HG power via Ephys and intrinsic signal via ISOI in response to left-eye or right-eye stimulation (160 trials each). The left and right panels show the responses of electrodes with left-eye or right-eye dominance, respectively. (g-h) Normalized responses of Ephys and ISOI before and after stroke at an electrode and the neighboring five pixels in the stroke region (g) and away from the stroke region (h); 160 trials per condition.

To investigate how these color or ocular-specific responses are correlated with the neural responses measured by the Smart Dura, we first selected two electrodes that strongly responded to chromatic or achromatic stimuli, respectively. We then quantified the averaged HG power for each electrode and the averaged time course of the intrinsic signals at five pixels near that electrode. The left panel in Figure 4e presents the HG power of the electrode that showed a chromatic dominance throughout the visual stimulation, peaking approximately 0.2-0.3 seconds after stimulus onset. The intrinsic responses exhibited slower dynamics with a peak delayed by about 3 seconds, but the chromatic dominance in the intrinsic signal was consistent with that in the HG power. A similar peak delay in the intrinsic signals, occurring a few seconds after the onset of the visual stimulus, has been reported in a previous study.^[41]^ In contrast, the right panel in Figure 4e shows results from an electrode having an achromatic dominance in HG power in the latter half of the visual stimulus, and a similar trend was observed in the intrinsic signals. To further examine the consistency between signals from electrical recording and optical imaging, we chose two electrodes that were highly activated during either left or right eye stimulation. As shown in Figure 4f, comparable ocular dominances were found between the HG power and the intrinsic signal. Lastly, we examined the evoked responses to visual stimulation before and after stroke. As described above, we induced an ischemic stroke in V1 via a photothrombotic lesioning technique and confirmed that the gamma power of the spontaneous activity in the stroke-induced region significantly decreased after the stroke. Here, we evaluated the changes in the visual stimulus-evoked HG power and intrinsic signal response caused by stroke induction. Figure 4g shows pre- and post-stroke responses at an electrode and five neighboring pixels within the stroke region, and both responses were strongly diminished after the stroke was induced. On the other hand, in the electrode far from the stroke region (Figure 4h), stimulus-locked increases in the HG power and intrinsic signal in response to visual stimulation remained strong. Collectively, the observed consistency between two modalities supports that the micron-sized electrodes of Smart Dura not only facilitate optical access to the areas surrounding the electrodes, but also allow for high-fidelity readout of spatially localized neural activity. Building on these findings, Smart Dura presents a great potential for integration with a variety of functional imaging techniques, thereby obtaining large-scale multimodal information for detailed analysis of spatiotemporal activity patterns in the brain.

### 2.5. Optogenetic Neuromodulation through Smart Dura

Simultaneous optogenetic neuromodulation and high-resolution electrophysiology enable selective manipulation of neural circuits while concurrently observing their impact on neural activity. Our Smart Dura offers spatially resolved electrophysiological recordings without compromising the optical access required for light delivery, facilitating precise optogenetic manipulation and evoked response characterization. To demonstrate the applicability of our Smart Dura for optogenetic experiments, we performed optogenetic modulation in an awake animal that had widespread expression of the inhibitory opsin Jaws. We infused a red-light-activated Jaws into the posterior parietal cortex (PPC) using convection-enhanced delivery, which provides large-scale viral vector delivery and opsin expression.^[25,42,43]^ At around two years after the initial injection, we conducted epifluorescence imaging through the Smart Dura, which revealed distinct opsin/GFP expression at two separate regions over the PPC (green regions in **Figure 5a**), similar to our previous observations.^[44]^ This shows the feasibility of epifluorescence monitoring of large-scale optogenetic expression through the Smart Dura.

**Figure 5.**
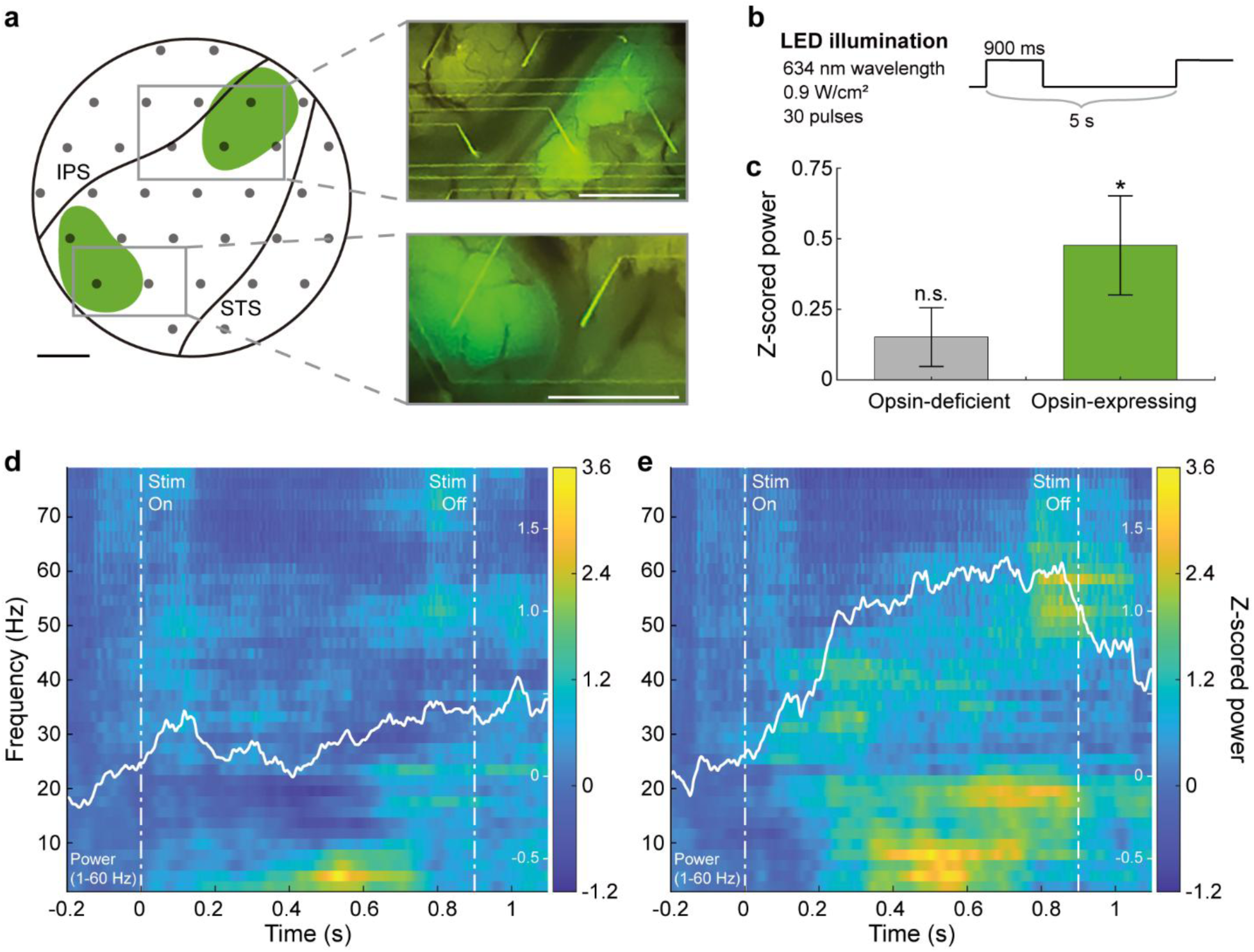
Characterization of evoked neural response during optogenetic modulation. (a) Optogenetic expression of opsin Jaws imaged through the Smart Dura over the PPC of the awake animal (IPS: intraparietal sulcus. STS: superior temporal sulcus). Green regions indicate locations with strong opsin/GFP expression. Insets on the right show optical access and epifluorescence over the two expressing regions at the start of optogenetic experiments. Scale bar: 2.5 mm. (b) Light stimulation pulses. (c) Light-evoked neural responses at individual electrodes in the opsin-deficient region and the opsin-expressing region. The broadband LFP (1-150 Hz) power during the stimulation significantly increased at the opsin-expressing location (p = 0.0109), whereas the power at the opsin-deficient location was not (p = 0.1548). Data are presented as the mean ± standard error of the mean. One-sample t-test. n = 30 trials. (d-e) Trial-averaged spectrograms of neural activity during light stimulation at electrodes in the opsin-deficient region (d) and opsin-expressing region (e). The light pulse onset occurred at t = 0 and lasted 900 ms. For each spectrogram, the averaged power in the 1-60 Hz frequency band is overlaid in white.

To activate the opsin, we delivered red-light stimuli (Figure 5b; 634 nm; 0.9 W/cm^2^) through the Smart Dura using a custom-built LED array^[44,45]^ to the entire cortical area. We then recorded neural activity and evaluated power changes in broadband local field potentials (LFPs) at frequencies of 1-150 Hz for individual electrodes in the opsin-expressing and opsin-deficient regions (Figure 5c). During the light stimulation, the broadband LFP power significantly increased in the area expressing the opsin (p = 0.0109; one-sample t-test) but not in the opsin-deficient region (p = 0.1548; one-sample t-test). A time-frequency representation of this differential response between opsin-deficient (Figure 5d) and opsin-expressing (Figure 5e) regions shows a significant power increase in the 1-60 Hz frequency band for the opsin-expressing region. This localized power increase, selective for opsin-expressing regions, is consistent with findings elaborated on in our previously published work^[44]^ and confirms the ability of the Smart Dura combined with optogenetics to modulate neural activity and simultaneously record frequency-specific evoked responses.

## 3. Discussion

In this study, we have demonstrated that Smart Dura functions as a unique multimodal neural interface for NHPs, enabling high-resolution electrophysiology, electrical stimulation, and optical access to the cortex. This capability unlocks great potential for multimodal and multiscale interrogation of brain function by bridging localized cellular activity with large-scale circuit dynamics. Through carefully designed experiments, we showed that Smart Dura is compatible with high-resolution optical readouts, including wide-field imaging such as OCTA and ISOI, as well as multiphoton imaging that enables single-cell resolution acquisition. Moreover, we established that Smart Dura facilitates integration with optical interventions, particularly optogenetic neuromodulation for cell-type-specific manipulations and photothrombotic lesioning for inducing focal ischemic stroke, positioning it as a powerful platform for studying both neural dynamics and causal mechanisms.

Our Smart Dura provides a large optical window with high transparency, as well as large-coverage, high-resolution electrophysiology. Using OCTA and ISOI, we were able to image blood flow (Figure 3b) and intrinsic signals (Figure 4c, d) across the entire 20 mm optical window. In this coverage area, the electrodes and metal traces with a width of tens of micrometers had negligible light-blocking effects compared to those in previous studies involving devices with hundreds of micrometers of opaque metal arrays.^[46,47]^ With multiphoton imaging, although the field of view was smaller than the other two imaging techniques, we obtained fluorescence images down to 550 μm with single-cell resolution, even beneath 10 μm-wide opaque metal traces (Figure 2). Furthermore, through light illumination at the millimeter scale, we performed targeted optical manipulations without interference from the electrode placement and captured the corresponding evoked responses – a decrease in gamma power in the stroke-induced region (Figure 3d) and an increase in broadband power in the opsin-expressing region following optogenetic stimulation (Figure 5e). Beyond demonstrating compatibility with individual techniques, an important strength of Smart Dura is its versatility in integrating multiple functionalities in a flexible, experiment-driven manner. Depending on the scientific question, electrical and optical methods can be combined in various ways to probe different aspects of brain function. For example, applying electrical stimulation together with electrophysiological recording and focal lesion induction allows for high-resolution investigation of the therapeutic effect of the stimulation following the acute ischemic stroke (Figure 3e). Likewise, integrating electrophysiology, ISOI, and photothrombotic lesioning enables assessment of functional deficits after stroke at multiple spatial and temporal scales (Figure 4g, h).

In addition to the demonstrations presented in this study, our Smart Dura holds great potential for technological improvements and new applications for future research. From a technological point of view, our microfabrication process can allow a further reduction of the opaque trace width from 50 μm, which caused shadowing in multiphoton imaging, to a negligible 10 μm or even narrower, minimizing light loss in optical technologies.^[21]^ Moreover, this scaling down facilitates the design of higher-density electrode arrays, improving the spatial resolution of electrophysiology and enabling more localized measurements of neural activity. Additionally, replacing the electrodes and traces with transparent conductive materials, such as graphene and indium tin oxide, can achieve further optical transparency.

From an application perspective, although we targeted in vivo multiphoton microscopy at fluorescent vasculature, functional neural imaging using calcium indicators or voltage-sensitive dyes can equally well be achieved through our Smart Dura. A few attempts at multiphoton calcium imaging for mapping at depth in the visual cortex of NHPs have been reported.^[48,49]^ The combination of electrophysiological and optical recording uniquely allows us to measure neuronal activity at different spatiotemporal scales, ranging from individual neurons to entire networks and from slow calcium signals to fast action potentials. This also provides an opportunity to obtain functional maps over the cortical surface and across cortical laminae. Furthermore, recent all-optical techniques that combine optical stimulation and readout have been proposed and demonstrated in NHPs.^[50,51]^ Integrating the electrical recording and stimulation capabilities of our Smart Dura with such optical functionality offers neural recording and modulation at multiple spatiotemporal scales, bringing us new insight into the complementary natures of multiscale information. This enables a comprehensive understanding of brain functions and innovative therapeutic paradigms for neurological disorders, ultimately being applicable and translatable to humans.

## 4. Experimental Section

### Animal Subjects, Surgical Procedures, and Electrophysiological Recording

Animal care and experiments were approved by the University of Washington’s Office of Animal Welfare, the Institutional Animal Care and Use Committee, and the Washington National Primate Research Center (WaNPRC). We tested our Smart Dura on two male macaques: *Macaca nemestrina* (4.8 kg, 4 years) was used for the experiments under anesthesia, and *Macaca mulatta* (12.6 kg, 11 years) was used for the awake experiments. The first animal underwent unilateral craniotomy and durotomy with a 25 mm diameter targeting the primary visual cortex (V1) in the left hemisphere. The animal was anesthetized with a continuous infusion of sufentanil citrate (4-6 μg/kg/hr) and vecuronium bromide (initial bolus of 0.1 mg/kg; 0.1 mg/kg/hr) throughout the experiment. Artificial respiration was adjusted to keep the expiratory carbon dioxide concentration within 3.8-4.0%, and body temperature was maintained at approximately 37°C using a heating pad. The second animal has a pre-established cranial chamber overlying the left posterior parietal cortex (PPC).^[44,45]^

For both animals, electrophysiological recordings were performed using a 64-channel Smart Dura, developed in our previous work.^[20,21]^ The electrodes and traces were fabricated with a Pt/Au/Pt metal stack on a PDMS and Parylene C bilayer substrate. The electrode size and spacing were 20 μm and 1818 μm, respectively, and PEDOT:PSS was electroplated on the electrodes to reduce the impedance. The details of the microfabrication process are described here.^[21]^ The Smart Dura was placed on the surface of the cortex and connected to the Front ends (Nano2+Stim; band-pass filter: 1 Hz-7.5 kHz), interfacing with a Grapevine Nomad processor and Trellis software (Ripple Neuro, UT). All recording components were supported by a custom 3D-printed holder that was affixed to the skull with dental acrylic for the anesthetized animal, with the grounding pins connected to a skull screw that was positioned anterior to the craniotomy. For the awake animal, the custom 3D-printed holder was screwed to the cranial chamber, and the ground pins were connected to the screw secured onto the chamber. Raw signals were digitized at a sampling rate of 30 kHz. Local field potentials (LFPs) were sampled at 1 kHz by low-pass filtering at 250 Hz with notch filters at multiples of 60 Hz. Subsequent signal processing and data analyses were carried out using MATLAB (MathWorks Inc., MA). Bad channels with high impedance and large noise were excluded from the analysis.

### High-spatial-resolution Transmission Measurement

We performed UV-Vis-IR spectroscopy using the optical setup shown in Figure 1c. With an incandescent light bulb as our broadband light source, a 100 μm diameter pinhole (P100HK; Thorlabs, NJ) and an iris (SM1D12D; Thorlabs) with its smallest aperture (0.8 mm in diameter) were used for generating 27 μm spot size and 167 μm spot size, respectively. The light beam passing through the pinhole or iris was collimated by an achromatic lens (f = 200 mm) (AC254-200-AB; Thorlabs) and then focused by another achromatic lens (f = 50 mm) (AC127-050-AB-ML; Thorlabs). A beam splitter (CM1-BS013; Thorlabs) was used to redirect the light path at a 90-degree angle onto our sample while enabling simultaneous imaging via a zoom lens (#88-399; Edmund Optics, NJ) and charged-coupled device (CCD) camera (BFS-U3-17S7C-C; FLIR, OR). An optical fiber (200 µm core diameter glass fiber; QP200-2-SR-BI) coupled to an optical spectrometer (HR4000CG-UV-VIS; Ocean Optics, FL) was placed directly under the Smart Dura to measure the transmission spectrum. Prior to placing the Smart Dura on top of the holder, we obtained measurements with the light source turned off (dark reading) as well as turned on (reference reading). The dark reading and reference reading were used to offset the effects of any coupled light into the fiber from the room environment and to obtain a baseline measurement without Smart Dura, respectively. With all three measurements: dark, reference, and sampling readings, the optical transmission of Smart Dura was determined by first subtracting the dark reading from both reference and sample readings, taking the ratio between the sample and reference, and then normalizing to the average value of the least blockage. For two light spot sizes of 27 and 167 μm in diameter, we measured transmission through the Smart Dura at different locations around the metal trace by 1D scanning (Figure S1).

### Multiphoton Imaging

Two- and three-photon imaging were conducted with a multi-photon microscope (MOM; Sutter Instruments, CA) equipped with excitation lasers set to 920 nm (Chameleon Vision II; Coherent, PA) and 1300 nm (Monaco/Opera-F; Coherent) wavelengths, respectively.^[52]^ The laser beam was focused with a 16x (CFI75 LWD 16X W; NA = 0.8; Nikon, Japan) or 25x (XLPLN25XWMP2; NA = 1.05; Olympus, Japan) water immersion objective lens. The microscope was controlled by ScanImage software (Vidrio Technologies, VA). For imaging of pollen grains, a microscope slide with mixed pollen grains (B690; Carolina Biological, NC) was utilized and the Smart Dura was located between the objective lens and the sample slide. The 3D reconstructed image in Figure 2b was acquired over a 260 μm thickness with 1 μm step increments. For in vivo vasculature imaging, the Smart Dura was placed on the V1 surface of the anesthetized animal and a custom 3D-printed well with an 18 mm diameter glass bottom (No. 1 thickness) was secured onto the Smart Dura to immerse the objective lens and flatten the area for imaging. FITC-dextran (50 mg/kg; 50 mg/mL in sterile water; 52471; Sigma-Aldrich, MA) was then intravenously injected. A depth of 0 µm for all images is set to the plane focused on the electrodes and metal traces. 3D image stacking, pseudo-coloring, image stitching, and intensity profile measurements were performed using ImageJ (National Institutes of Health, MD).

### Photothrombotic Lesioning followed by Electrical Stimulation

We induced photochemical ischemic lesioning in V1 using the photothrombotic technique.^[31]^ The Smart Dura was placed on the cortical surface and an opaque silicone mask with a 1.5 mm-diameter circular aperture in the center was placed over it. A photoactive dye, Rose Bengal (20 mg/kg; 40 mg/mL in saline; excitation peak at 559 nm; 330000; Sigma-Aldrich), was infused intravenously for 5 minutes simultaneously with the start of illumination using a cold white light source (KL 2500 LCD; SCHOTT, UK). Light illumination through the mask aperture lasted for 30 minutes.

We recorded pre-stroke and post-stroke neural activity for 30 and 60 minutes, respectively, and then delivered theta-burst electrical stimuli to an electrode approximately 6.5 mm medial to the lesion center. Electrical stimulation was applied with the same parameters as those used in our previous work,^[33]^ except that the stimulation amplitude was 30 μA. Between each stimulation block, 2 minutes of spontaneous activity was recorded. The averaged gamma power in the frequency band of 30-59 Hz for each electrode was calculated by squaring the signal and dividing it by its duration. To compare gamma power, we used a repeated measures ANOVA with Bonferroni’s post-hoc test. Pairwise comparisons were only performed between pre-stroke and other recording blocks and between post-stroke and post-electrical stimulation recording blocks.

### Optical Coherence Tomography Angiography

We used a custom-built optical coherence tomography (OCT) system for OCT angiography (OCTA) imaging. The prototype is similar to the device, reported in our previous publications.^[36,37]^ Briefly, the system employed a swept-source laser operating at 200 kHz, with a center wavelength of 1310 nm and an optical bandwidth of 100 nm, providing an axial resolution of about 8 µm in tissue. The objective lens (LSM04; Thorlabs) had a focal length of 54 mm and provided a lateral resolution of about 35 µm. The total beam power incident to the Smart Dura was 5 mW. The acquisition time of one OCT data volume took 8 seconds. Each data volume covered a 9×9 mm^2^ cortical area, rasterized into 500 cross-sectional planes with each plan containing 500 A-scans. Four repeated B-scans were acquired at every individual plane for contrasting blood flow. During the imaging session, five data volumes were consecutively acquired at the center, anterior-medial, anterior-lateral, posterior-lateral, and posterior-medial sections of the cranial window to cover the entire area (i.e., the center and four corners). The system-embedded RGB camera was used to navigate the imaging probe in real-time. The OCTA image processing was conducted in MATLAB. An eigen-decomposition-based algorithm^[53]^ was first used to contrast the blood flow signal. Then, top-down max-intensity projection was performed to suppress the blood flow volumes onto two-dimensional vascular maps, with pseudo-color encoding to indicate the depth of the vessels. Finally, these individual maps were stitched together to form a larger map covering the entire region.

### Visual Stimulation

We presented chromatic and achromatic square drifting grating (4 s, 1.0 cycles/degree, 4.0 degree/s) on the display (Display++; Cambridge Research Systems) at a distance of 76 or 63 cm. The chromatic drifting grating consisted of red and green in the same luminance (17.4 cd/m^2^). The achromatic stimulus consisted of white and black (34.6 and 0.12 cd/m^2^, respectively), so the mean luminance was adjusted to the same level as that of the chromatic stimulus. Eight directions were randomly presented in one session for 10 repeats for each condition. One eye was occluded with a shield plate for each session. To characterize the neural responses evoked by visual stimuli from electrophysiological recordings, a multitaper spectrogram was generated in a window of 300 ms with 3-ms steps for each trial. Each time-frequency bin was z-scored with respect to the frequency band power 4 seconds before the onset of stimulation. The z-scored spectrograms were averaged across trials, and high-gamma (HG) power was obtained from bins 60-150 Hz of the trial-averaged spectrogram.

### Wide-field Intrinsic Signal Optical Imaging

The image sequences were acquired with a sCMOS camera (Zyla-4.2 USB3; Andor) equipped with a tandem lens configuration (Nikkor 50 mm and Nikkor 85 mm; Nikon) to achieve a 22.6 mm field of view. The cortex was illuminated with an LED at 625 nm wavelength (M625L4; Thorlabs) through the Smart Dura. Image acquisition was controlled by Micro-manager software.^[54]^ The response amplitude was calculated as a fraction of the image intensity from the baseline. First, baseline intensity R_0_ was calculated as an average over one second before the stimulus onset. Then, R(t) was calculated as an average with a shifting time window of a one-second width. Then, the amplitude was calculated as ΔR(t)/R_0_ = (R(t)-R_0_)/R_0_. The color preference pattern was visualized as a t-statistic between the chromatic and achromatic conditions.

The ocular dominance pattern was similarly visualized by comparing the left eye and the right eye conditions. The vasculature pattern was also acquired with the same imaging setup under 530 nm illumination (M530L4; Thorlabs) for later alignment of Smart Dura contact positions. For the comparison with HG power measured by the Smart Dura, time courses of the intrinsic responses were obtained at five pixels near each selected electrode and averaged.

### Optogenetic Neuromodulation

During the surgical implantation of a chronic chamber as described in our previous works,^[44,45]^ we virally infused a red-light-activated inhibitory opsin Jaws^[55]^ into the PPC using convection-enhanced delivery^[25]^ of viral vector rAAV8-hSyn-Jaws-KGC-GFP-ER2 (UNC Vector Core, NC). The optogenetic experiment demonstrated in this work was performed around 2 years after the initial injection. To test the expression level at different locations over the optical window, we performed epifluorescence imaging of the GFP fluorescence tag through the Smart Dura after the dura resection using a blue light source (440-460 nm excitation; SFA-RB; NIGHTSEA, PA) and a green emission filter (500-560 nm bandpass) attached to a digital camera with 35mm lens (D5300; Nikon). Only for the epifluorescence imaging, we used a 32-channel Smart Dura. Each quadrant of the optical window was illuminated sequentially during imaging, and the images were stacked and averaged in Photoshop (Adobe, CA) to improve quality and contrast over the full window.

To activate the opsin by applying light stimulation, an LED array was placed over the Smart Dura, as described in detail in our previous publications.^[44,45]^ The array consisted of a 4×4 grid of LEDs emitting light at a wavelength of 634 nm at 0.9 W/cm^2^ over the entire cortical area (about 1.0 cm^2^). 30 light pulses were delivered every 5 seconds, lasting 900 ms. To remove the photoelectric artifact generated by the light pulse, an electrode recording in saline was subject to the same light stimulation paradigm, and the first three principal components (PCs) of the artifact during the 900 ms of stimulation and 900 ms after stimulation were used to generate two templates of the artifact without neural activity, respectively. Each stimulation pulse in the neural recording was segmented into 900 ms during and 900 ms after stimulation and then projected onto the PC space of the corresponding saline template. The scaled projection of the neural data was then subtracted from the original neural recording to remove the artifact. To characterize the light-evoked neural response, a multitaper spectrogram with a window of 300 ms with 3-ms steps for each trial was used. Each time-frequency bin was z-scored with respect to the frequency band power 350 ms before the onset of stimulation. The time-frequency bins during the stimulation were then averaged between 1-150 Hz to calculate the broadband LPF power change. The evoked response was considered significant if the z-scored power during stimulation was significantly greater using a one-sample t-test, where the null hypothesis is rejected when the mean is not equal to 0.

## Acknowledgments

This work was supported by the National Institute of Neurological Disorders and Stroke of the National Institutes of Health (NIH) R01NS116464, R01NS119395, and U01NS115585, the National Institute of Mental Health of the NIH R01MH125429 and UG3MH126864, the Washington National Primate Research Center P51OD010425 and U42OD011123 from the NIH Office of Research Infrastructure Programs, the Weill Neurohub, the Washington Research Foundation, the American Heart Association, the NIH T32 training in theoretical and computational approaches to neural circuits of cognition 1T32MH132518, the National Institute of Biomedical Imaging and Bioengineering of the NIH T32EB029365, and the Bertucci Nanotechnology Laboratory at Carnegie Mellon University BNL-78657879. We thank Toni Haun, Celeste Dylla, Anne F. Pierce, Felix Schwock, Sophia Shan, and WaNPRC veterinarians and staff for their help with animal care, experimental preparation, and assistance during surgeries.

## References

[1] A. Vázquez-Guardado, Y. Yang, A. J. Bandodkar, J. A. Rogers, Nat. Neurosci. 2020 2312 2020, 23, 1522.

[2] E. S. Boyden, F. Zhang, E. Bamberg, G. Nagel, K. Deisseroth, Nat. Neurosci. 2005 89 2005, 8, 1263.

[3] Z. Fekete, A. Zátonyi, A. Kaszás, M. Madarász, A. Slézia, Microsystems Nanoeng. 2023 91 2023, 9, 1.

[4] Y. U. Cho, S. L. Lim, J. H. Hong, K. J. Yu, npj Flex. Electron. 2022 61 2022, 6, 1.

[5] M. Ramezani, Y. Ren, E. Cubukcu, D. Kuzum, Nat. Rev. Electr. Eng. 2024 21 2024, 2, 42.

[6] X. Liu, C. Ren, Y. Lu, Y. Liu, J. H. Kim, S. Leutgeb, T. Komiyama, D. Kuzum, Nat. Neurosci. 2021 246 2021, 24, 886.

[7] M. Yuan, F. Li, F. Xue, Y. Wang, B. Li, R. Tang, Y. Wang, G. Q. Bi, W. Pei, Microsystems Nanoeng. 2025 111 2025, 11, 1.

[8] A. F. Renz, J. Lee, K. Tybrandt, M. Brzezinski, D. A. Lorenzo, M. Cerra Cheraka, J. Lee, F. Helmchen, J. Vörös, C. M. Lewis, Adv. Healthc. Mater. 2020, 9, 2000814.

[9] M. Thunemann, Y. Lu, X. Liu, K. Klllç, M. Desjardins, M. Vandenberghe, S. Sadegh, P. A. Saisan, Q. Cheng, K. L. Weldy, H. Lyu, S. Djurovic, O. A. Andreassen, A. M. Dale, A. Devor, D. Kuzum, Nat. Commun. 2018 91 2018, 9, 1.

[10] J. Zhang, X. Liu, W. Xu, W. Luo, M. Li, F. Chu, L. Xu, A. Cao, J. Guan, S. Tang, X. Duan, Nano Lett. 2018, 18, 2903.

[11] M. J. Donahue, A. Kaszas, G. F. Turi, B. Rózsa, A. Slézia, I. Vanzetta, G. Katona, C. Bernard, G. G. Malliaras, A. Williamson, eNeuro 2018, 5.

[12] N. Kunori, I. Takashima, J. Neurosci. Methods 2015, 251, 130.

[13] A. Zátonyi, Z. Borhegyi, M. Srivastava, D. Cserpán, Z. Somogyvári, Z. Kisvárday, Z. Fekete, Sensors Actuators B Chem. 2018, 273, 519.

[14] T. J. Richner, S. Thongpang, S. K. Brodnick, A. A. Schendel, R. W. Falk, L. A. Krugner-Higby, R. Pashaie, J. C. Williams, J. Neural Eng. 2014, 11, 016010.

[15] D. W. Park, A. A. Schendel, S. Mikael, S. K. Brodnick, T. J. Richner, J. P. Ness, M. R. Hayat, F. Atry, S. T. Frye, R. Pashaie, S. Thongpang, Z. Ma, J. C. Williams, Nat. Commun. 2014 51 2014, 5, 1.

[16] Y. U. Cho, J. Y. Lee, U. J. Jeong, S. H. Park, S. L. Lim, K. Y. Kim, J. W. Jang, J. H. Park, H. W. Kim, H. Shin, H. Jeon, Y. M. Jung, I. J. Cho, K. J. Yu, Adv. Funct. Mater. 2022, 32, 2105568.

[17] D. Kim, M. Bissannagari, B. Kim, N. Hong, J. Park, H. Lim, J. Lee, J. Lee, Y. K. Kim, Y. Cho, K. Lee, J. Lee, J.-H. Yoon, J. E. Jang, D. Tsai, S. Lee, H.-J. Kwon, H. K. Choe, H. Kang, npj Flex. Electron. 2025 91 2025, 9, 1.

[18] D. J. O’Shea, E. Trautmann, C. Chandrasekaran, S. Stavisky, J. C. Kao, M. Sahani, S. Ryu, K. Deisseroth, K. V. Shenoy, Exp. Neurol. 2017, 287, 437.

[19] T. Belloir, S. Montalgo-Vargo, Z. Ahmed, D. J. Griggs, S. Fisher, T. Brown, M. Chamanzar, A. Yazdan-Shahmorad, iScience 2023, 26, 105866.

[20] S. M. Vargo, T. Belloir, I. Kimukin, Z. Ahmed, D. J. Griggs, N. Stanis, A. Yazdan-Shahmorad, M. Chamanzar, Int. IEEE/EMBS Conf. Neural Eng. NER 2023, 2023-April.

[21] S. M. Vargo, N. Hong, T. Belloir, N. Stanis, J. Zhou, K. Khateeb, G. Hatanaka, Z. Ahmed, I. Kimukin, D. J. Griggs, W. Bair, A. Yazdan-Shahmorad, M. Chamanzar, bioRxiv 2025, 2025.02.26.640369.

[22] A. Arieli, A. Grinvald, H. Slovin, J. Neurosci. Methods 2002, 114, 119.

[23] H. Slovin, A. Arieli, R. Hildesheim, A. Grinvald, J. Neurophysiol. 2002, 88, 3421.

[24] O. Ruiz, B. R. Lustig, J. J. Nassi, A. Cetin, J. H. Reynolds, T. D. Albright, E. M. Callaway, G. R. Stoner, A. W. Roe, J. Neurophysiol. 2013, 110, 1455.

[25] A. Yazdan-Shahmorad, C. Diaz-Botia, T. L. Hanson, V. Kharazia, P. Ledochowitsch, M. M. Maharbiz, P. N. Sabes, Neuron 2016, 89, 927.

[26] D. Khodagholy, J. N. Gelinas, T. Thesen, W. Doyle, O. Devinsky, G. G. Malliaras, G. Buzsáki, Nat. Neurosci. 2015 182 2014, 18, 310.

[27] C. Xu, M. Nedergaard, D. J. Fowell, P. Friedl, N. Ji, Cell 2024, 187, 4458.

[28] T. H. Kim, M. J. Schnitzer, Cell 2022, 185, 9.

[29] N. G. Horton, K. Wang, D. Kobat, C. G. Clark, F. W. Wise, C. B. Schaffer, C. Xu, Nat. Photonics 2013 73 2013, 7, 205.

[30] D. G. Ouzounov, T. Wang, M. Wang, D. D. Feng, N. G. Horton, J. C. Cruz-Hernández, Y. T. Cheng, J. Reimer, A. S. Tolias, N. Nishimura, C. Xu, Nat. Methods 2017 144 2017, 14, 388.

[31] K. Khateeb, J. Bloch, J. Zhou, M. Rahimi, D. J. Griggs, V. N. Kharazia, M. N. Le, R. K. Wang, A. Yazdan-Shahmorad, Cell Reports Methods 2022, 2, 100183.

[32] K. Khateeb, Z. Yao, V. N. Kharazia, E. P. Burunova, S. Song, R. Wang, A. Yazdan-Shahmorad, Proc. Annu. Int. Conf. IEEE Eng. Med. Biol. Soc. EMBS 2019, 3515.

[33] J. Zhou, K. Khateeb, A. Yazdan-Shahmorad, Nat. Commun. 2025 161 2025, 16, 1.

[34] J. Zhou, K. Khateeb, A. Gala, M. Rahimi, D. J. Griggs, Z. Ip, A. Yazdan-Shahmorad, Proc. Annu. Int. Conf. IEEE Eng. Med. Biol. Soc. EMBS 2022, 2022-July, 3085.

[35] N. Stanis, K. Khateeb, J. Zhou, R. K. Wang, A. Yazdan-Shahmorad, STAR Protoc. 2023, 4, 102496.

[36] Z. Xie, Y. Shi, A. Marmin, R. K. Wang, J. Biophotonics 2024, e202400289.

[37] A. J. Deegan, J. Lu, R. Sharma, S. P. Mandell, R. K. Wang, Quant. Imaging Med. Surg. 2021, 11, 78496.

[38] A. Grinvald, E. Lieke, R. D. Frostig, C. D. Gilbert, T. N. Wiesel, Nat. 1986 3246095 1986, 324, 361.

[39] J. C. Horton, D. H. Hubel, Nat. 1981 2925825 1981, 292, 762.

[40] D. H. Hubel, T. N. Wiesel, Nat. 1969 2215182 1969, 221, 747.

[41] H. D. Lu, G. Chen, J. Cai, A. W. Roe, Neuroimage 2017, 148, 160.

[42] K. Khateeb, D. J. Griggs, P. N. Sabes, A. Yazdan-Shahmorad, J. Vis. Exp. 2019.

[43] D. J. Griggs, A. D. Garcia, W. Y. Au, W. K. S. Ojemann, A. G. Johnson, J. T. Ting, E. A. Buffalo, A. Yazdan-Shahmorad, Pharmaceutics 2022, 14, 1435.

[44] D. J. Griggs, J. Bloch, N. Stanis, J. Zhou, S. Fisher, H. Jahanian, A. Yazdan-Shahmorad, bioRxiv 2024, 2024.06.25.600719.

[45] D. J. Griggs, J. Bloch, S. Fisher, W. K. S. Ojemann, K. M. Coubrough, K. Khateeb, M. Chu, A. Yazdan-Shahmorad, Proc. Annu. Int. Conf. IEEE Eng. Med. Biol. Soc. EMBS 2022, 2022-July, 3081.

[46] D. W. Park, S. K. Brodnick, J. P. Ness, F. Atry, L. Krugner-Higby, A. Sandberg, S. Mikael, T. J. Richner, J. Novello, H. Kim, D. H. Baek, J. Bong, S. T. Frye, S. Thongpang, K. I. Swanson, W. Lake, R. Pashaie, J. C. Williams, Z. Ma, Nat. Protoc. 2016 1111 2016, 11, 2201.

[47] D. J. Griggs, K. Khateeb, J. Zhou, T. Liu, R. Wang, A. Yazdan-Shahmorad, J. Neural Eng. 2021, 18, 055006.

[48] I. Nauhaus, K. J. Nielsen, A. A. Disney, E. M. Callaway, Nat. Neurosci. 2012 1512 2012, 15, 1683.

[49] S. Chatterjee, K. Ohki, R. C. Reid, Nat. Commun. 2021 121 2021, 12, 1.

[50] Y. Nakamichi, K. Okubo, T. Sato, M. Hashimoto, M. Tanifuji, Sci. Reports 2019 91 2019, 9, 1.

[51] N. Ju, R. Jiang, S. L. Macknik, S. Martinez-Conde, S. Tang, PLOS Biol. 2018, 16, e2005839.

[52] K. Takasaki, R. Abbasi-Asl, J. Waters, eNeuro 2020, 7.

[53] S. Yousefi, Z. Zhi, R. K. Wang, IEEE Trans. Biomed. Eng. 2011, 58, 2316.

[54] A. Edelstein, N. Amodaj, K. Hoover, R. Vale, N. Stuurman, Curr. Protoc. Mol. Biol. 2010, 92, 14.20.1.

[55] A. S. Chuong, M. L. Miri, V. Busskamp, G. A. C. Matthews, L. C. Acker, A. T. Sørensen, A. Young, N. C. Klapoetke, M. A. Henninger, S. B. Kodandaramaiah, M. Ogawa, S. B. Ramanlal, R. C. Bandler, B. D. Allen, C. R. Forest, B. Y. Chow, X. Han, Y. Lin, K. M. Tye, B. Roska, J. A. Cardin, E. S. Boyden, Nat. Neurosci. 2014 178 2014, 17, 1123.

